# Fast Set Operations for Compact *k*-mer Sets

**DOI:** 10.64898/2026.05.24.727514

**Authors:** Jarno N. Alanko, Lore Depuydt, Camille Marchet, Simon J. Puglisi

## Abstract

The *k*-mer spectrum of a set of sequences is the set of *k*-length substrings the sequences contain. This lossy representation of sequence content pervades modern genomics. Recently, the spectral Burrows-Wheeler transform (SBWT) has emerged as a space-efficient representation of *k*-spectra that also supports efficient *k*-mer lookup queries and, more generally, easy navigation of the de Bruijn graph of the *k*-spectrum. In this paper, we examine primitive set operations, such as intersection, union, and set difference, on SBWT-encoded *k*-spectra and show that these operations can be supported efficiently. Moreover, efficient merging leads directly to a new memory-efficient algorithm for SBWT construction, which was able to build the SBWT for the 661K bacterial dataset containing 88 billion distinct *k*-mers in 50 hours using 186 GiB of RAM and 112 GiB of disk space. Given the pervasiveness of *k*-mer sets in genomics and the continued rapid growth of genomic databases, our work opens the door to a wide array of future applications that manipulate and reason about genomic data by dealing directly with simultaneously compact and searchable *k*-mer set representations offered by the SBWT.

**2012 ACM Subject Classification:** Theory of computation → Design and analysis of algorithms

**Digital Object Identifier:** 10.4230/LIPIcs.WABI.2026.

**Supplementary Material:** *Software (Source Code)*: https://github.com/LoreDepuydt/sbwt-set-operations

**Funding:** This work has benefited from funding from the French State under the France 2030 program, reference ANR-21-IDES-0006. The European Metropolis of Lille and the University of Lille are also acknowledged for their funding and support of the project WILL-CHAIRES-25-001-BOSSA.

## 1 Introduction

The *k*-mer spectrum of a set of sequences is the set of *k*-length substrings that the sequences contain and is a lossy representation of sequence content that pervades modern genomics. *k*-mers are amenable to hashing, counting, and indexing at speeds that alignment-based approaches cannot match, and they are present in a wide range of applications from genome assembly to taxonomic classification. In rough terms, *k*-mers let pipelines scale by sacrificing an acceptable loss of fidelity (long-range relationships) for tractability.

This scalability is especially important given the recent emergence of peta-scale *k*-mer indexes over public sequence collections [7, 12], which allow researchers to query the entirety of deposited sequencing data in minutes. However, an open problem is that these systems are largely static, updates require costly rebuilds, and supported queries are narrow. As repositories such as AllTheBacteria [11] and the Human Pangenome Reference Consortium [19] regularly release new assemblies, freeze-and-rebuild will become unsustainable.

Beyond the index update problem, biologically meaningful questions—essentially questions of data mining—such as determining the set of *k*-mers shared across a panel of conditions [8], require boolean set operations over indexed *k*-mer sets. This type of query is not yet supported in large scale systems, as in-place set operations on compact *k*-mer representations remain both time- and space-intensive [14].

### Contribution

An ideal *k*-mer representation is simultaneously (i) compact, fitting large collections in modest hardware; (ii) searchable, supporting fast membership and navigation queries; and (iii) composable, allowing indexes to be combined via union, intersection, and difference without rebuilding from raw sequence. The first two are met by the spectral Burrows-Wheeler transform (SBWT) [5], which represents *k*-spectra space-efficiently while supporting *k*-mer lookup and de Bruijn graph navigation in time independent of set size.

In this paper, we address the third problem above by giving union, intersection, and set difference on SBWT-encoded *k*-spectra, built around a compact 𝒪 (*n*)-bit merge plan constructible in 𝒪 (*nk*) time. As a direct application, we build an SBWT index over the full 661K bacterial dataset [6], consisting of 2.57 Tbp and 88.70 billion distinct 31-mers, in 50 hours using 186 GiB peak RAM and 112 GiB peak disk space, where direct construction would need an estimated 1.2 TiB of RAM or 5.6 TiB of disk space.

### Related work

Set operations on *k*-mer collections have been approached from several angles. KMC [13] stores *k*-mers in sorted, partitioned databases and supports union, intersection, and subtraction via streaming merges; CBL [14] provides the same three operations on compact in-memory *k*-mer sets, and de Bruijn graph indexes such as Bifrost [10] and MetaGraph [12] expose merge operations on their respective representations. Minimal perfect hashing approaches (e.g., SSHash [17]) offer fast queries, but combining two indexes requires rebuilding the *k*-mer representation and hash function from scratch. On the indexed-Burrows-Wheeler side, Muggli et al. [16] give an O(*nk*)-time algorithm for merging BOSS representations of de Bruijn graphs, and Egidi et al. [9] describe an O(*n*^2^)-time algorithm for general Wheeler graphs together with a space-improved O(*nk*) variant for BOSS. Alanko et al. [1] support dynamic *k*-mer sets via an O(*nk*) merge on a Wheeler index of a de Bruijn graph, simulating deletions with a bit vector.

### Paper Outline

Section 2 fixes notation for *k*-spectra and the SBWT. Section 3.1 introduces the *merge plan*, a compact 𝒪 (*n*)-bit encoding of how two padded *k*-spectra interleave in colexicographic order, and gives an 𝒪 (*nk*)-time construction algorithm together with a sketch of an 𝒪 (*n*log*k*) variant. In Section 3.2, we use the merge plan to construct functional SBWT indexes of the union, intersection, and set difference. Section 4 evaluates the resulting algorithms against KMC and a sorted-list baseline, and applies the merge operation to build an SBWT over the full 661K bacterial dataset.

## 2 Preliminaries

Throughout this paper, a *string X*[1..*n*] is a sequence of *X* symbols over the DNA alphabet ∑ = {A,C,G,T}. The empty string is denoted ϵ and |ϵ| = 0. The *substring* of *X* starting at symbol *i* and ending at symbol *j* is denoted *X*[*i*..*j*]. We also use the half-open interval notations *X*(*i*..*j*] =*X*[*i*+1..*j*] and *X*[*i*..*j*) =*X*[*i*..*j* −1]. A *prefix* is a substring with starting position 1 and a *suffix* is a substring with ending position *n*.

A *k-mer* refers to a (sub)string of length *k*. The set of distinct *k*-mers of a string *X* is called the *k-spectrum* of *X*.

### Definition 1

*(k-Spectrum) The k-spectrum S*_*k*_(*X*) *of a string X is the set of distinct k-mers* {*X*[*i*..*i*+*k*−1] | *i*= 1,…,|*X*|−*k*+1}.

### Definition 2

*(k-Spectrum of a set of strings) The k-spectrum S*_*k*_(*X*_1_,…,*X*_*m*_) *of a set of strings X*_1_,…,*X*_*m*_ *is the union* 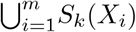.

A *padded k-spectrum* adds a minimal set of $-padded *dummy k*-mers to ensure that in the de Bruijn graph of the *k*-mers, every non-dummy *k*-mer has an incoming path of length at least *k*:

### Definition 3

*(Padded k-Spectrum) Let R*=*S*_*k*_(*X*_1_,…,*X*_*m*_) *be a k-spectrum with alphabet* ∑, *and let R*′⊆ *R be the set of-mers such that Y* [1..*k* − 1] *is not a suffix of any-mer in*. *The padded k-spectrum is the set* 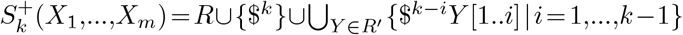, *with special character* $∉∑ *and* $<*c for all c*∈∑.

For example, if *X*_1_ = ACGT, *X*_2_ = GACG and *k* = 3, then *S*_3_(*X*_1_,*X*_2_) = {ACG,CGT,GAC} and 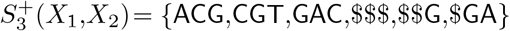. The *k*-mers that contain the dollar-symbols are called *dummy k-mers*.

The spectral Burrows-Wheeler transform maps a padded *k*-spectrum into a sequence of subsets of the alphabet:

### Definition 4

*(Spectral Burrows-Wheeler transform (SBWT)) Let R*^+^ *be a padded k-spectrum and let* 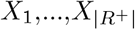 *be the elements of R*^+^ *in colexicographic order (i*.*e*., *comparing the k-mers from right to left). The SBWT is the sequence of sets* 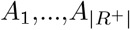 *with A*_*i*_ ⊆∑ *such that A*_*i*_ =∅ *if i*> 1 *and X*_*i*_[2..*k*] =*X*_*i*−1_[2..*k*], *otherwise A*_*i*_ ={*c*∈∑|*X*_*i*_[2..*k*]*c* ∈*R*^+^}.

It is helpful to think of the SBWT as encoding the set *R*^+^ of padded *k*-mers as a list arranged in colexicographic order. The *SBWT graph* is a subgraph of the node-centric de Bruijn graph of the *k*-mers, which contains only the smallest incoming de Bruijn graph edge into each node, where edges are compared by the colexicographic order of their origin nodes. This graph gives an alternative and equivalent way of defining the SBWT subset sequence: the *i*-th set in the SBWT sequence is the set of outgoing edge labels in the SBWT graph from the *i*-th *k*-mer in colexicographic order. We refer the reader to prior work [5, 3, 2] for examples and a more in-depth discussion of the SBWT.

In practice, the SBWT is represented as a bit matrix with one column col_*c*_ per character *c* ∈ ∑, where row *p* has a set bit in column col_*c*_ if *c* ∈ *A*_*p*_. From now on, we write SBWT[*p*] for the set *A*_*p*_. Let *C*[*c*] be the number of edge labels smaller than *c* in the SBWT graph. For a row *p* and character *c* ∈ SBWT[*p*], let succ(*p,c*) denote the destination node reached by following the outgoing edge labelled *c* from row *p*. This operation is supported by the default SBWT representation [5]: succ(*p,c*) = 1+*C*[*c*]+subsetrank_*c*_(*p*), where subsetrank_*c*_(*p*) is the number of rows *j* ≤ *p* such that *c* ∈ SBWT[*j*]. The +1 skips the source node corresponding to *k*-mer $^*k*^. Conversely, for a row *q* ≠ 1, let pred(*q*) denote the predecessor reached by following the unique incoming-character edge of *q*. This operation requires select support in the SBWT bit matrix, whose supporting data structures are not stored by default but can be added on demand. In the bit matrix representation, pred(*q*) =subsetselect_*c*_(*q*−1−*C*[*c*]), where *c* is the unique character satisfying 1+*C*[*c*] <*q* ≤ 1+*C*[*c*]+|col_*c*_|, and subsetselect_*c*_(*r*) returns the row containing the *r*-th set bit in column col_*c*_.

## 3 Set operations on SBWTs

### 3.1 Merge plan

Let *A*^+^ and *B*^+^ be two sets of *k*-mers (including the necessary dummy *k*-mers in their respective SBWTs). In order to support set operations, we first compute a *merge plan* of *A*^+^ and *B*^+^. Conceptually, this merge plan is a string *T* of length |*A*^+^ ∪ *B*^+^|, from the alphabet {L,R,B}, where the symbols stand for L = “left”, R = “right” and B “both”. The string is defined as follows:

- *T* [*i*] = L if the colexicographically *i*-th *k*-mer in *A*^+^ ∪ *B*^+^ is found only in *A*^+^
- *T* [*i*] = R if the colexicographically *i*-th *k*-mer in *A*^+^ ∪ *B*^+^ is found only in *B*^+^
- *T* [*i*] = B otherwise (found in both *A*^+^ and *B*^+^).

With *T* and the colexicographically sorted lists of *k*-mers in *A*^+^ and *B*^+^, we can list the *k*-mers of *A*^+^ ∪ *B*^+^ in colexicographic order by placing a pointer to the starts of *A*^+^ and *B*^+^, and picking *k*-mers from *A*^+^ and *B*^+^ according to *T* while advancing the pointers. This amounts to computing the union of *A*^+^ and *B*^+^.We can also perform an intersection by reporting only the *k*-mers that are associated with the letter B in *T*, or a set difference by reporting only *k*-mers associated with the letter L for *A*^+^\*B*^+^, or R for *B*^+^\*A*^+^.

#### 3.1.1 Computing the merge plan in *O*(*nk*) time and *O*(*n*log*σ*) bits

Let 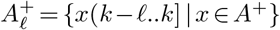 be the suffixes of length ℓ in *A*^+^, and similarly let 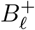 be the suffixes of length ℓ in *B*^+^. Let *P*_ℓ_ be the merge plan for 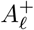 and 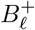. We represent *P*_ℓ_ as a sequence of pairs (*a*_0_,*b*_0_),…(*a*_*m*−1_,*b*_*m*−1_), where 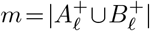. The number *a*_*i*_ is the number of *k*-mers in *A*^+^ that are suffixed by the *i*-th ℓ-mer of 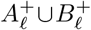, and the number *b*_*i*_ is the number of *k*-mers in *B*^+^ that are suffixed by the *i*-th *ℓ*-mer 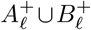. Some of these numbers can be zero, and we always have 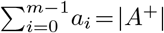 and 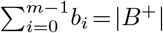.

We initialize the algorithm by setting *P*_0_ = (|*A*^+^|,| B^+^|),, and run *k* rounds to refine *P*_0_ to *P*_1_, then to *P*_2_, and so on up to *P*_*k*_. Let *M*_*X*_ be the *k-mer matrix* of a set of *k*-mers *X*, such that the rows of the matrix are the *k*-mers of *X* in colexicographic order. Now, given *P*_ℓ_,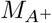 and 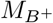, it is straightforward to stream over the pairs (*a*_*i*_,*b*_*i*_) in *P*_ℓ_, and output the pairs for *P*_ℓ+1_: We maintain the current range of *k*-mers both in 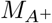 and 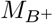, and refine each pair (*a*_*i*_,*b*_*i*_) into up to |∑| pieces based on the characters at the (ℓ+1)-th column from the right. After *k* rounds, the only pairs that can possibly remain in *P*_*k*_ are (1,0), (0,1) and (1,1). These correspond to the letters L, R and B in the definition of *T* above.

To save space, we encode the numbers *a*_*i*_ and *b*_*i*_, in unary such that integer *x* ≥ 0 is encoded rep(*x*) = 0^*x*^1. A list of pairs (*a*_0_,*b*_0_),(*a*_1_,*b*_1_)… is then encoded by concatenating the codes `rep`(*a*_0_), `rep`(*a*_1_) …, and concatenating the codes rep(*b*_0_), rep(*b*_1_) …. This takes a total of *O*(|*A*^+^ ∪ *B*^+^|) bits of space. With this, we give up the ability for random access in the list of pairs, but we are still able to stream the pairs in order, which is all we need.

The *k*-mer matrix *M*_*X*_ is never materialized in memory in full – instead, we materialize the matrix column by column from right to left using the SBWT. Algorithm 1 shows how the SBWT is used to transform one column of the *k*-mer matrix into the column on the left. For the first iteration, `column_in` is initialized to the last column of the *k*-mer matrix, which is obtained directly from the *C*-array. Similar algorithms can be found in the literature in the context of the BOSS data structure [16] and Wheeler graphs [1]. The memory access pattern of the algorithm is |∑| linearly advancing pointers, meaning that most of the computation is confined within the CPU cache, making the algorithm very fast in practice.

The algorithm is also cleanly parallelizable into independent threads by assigning disjoint input and output memory regions for each thread. The work can be split evenly for *t* output threads by splitting the input array `column_in` into *t* equal pieces, and using binary search on rank queries to find the ranges in `column_out` corresponding to each input piece.

##### Algorithm 1

Transforming a column in the SBWT *k*-mer matrix to the column on the left. In our implementation, the 1-bits in each bit matrix column are enumerated using leading-zero machine instructions to find the next set bit.

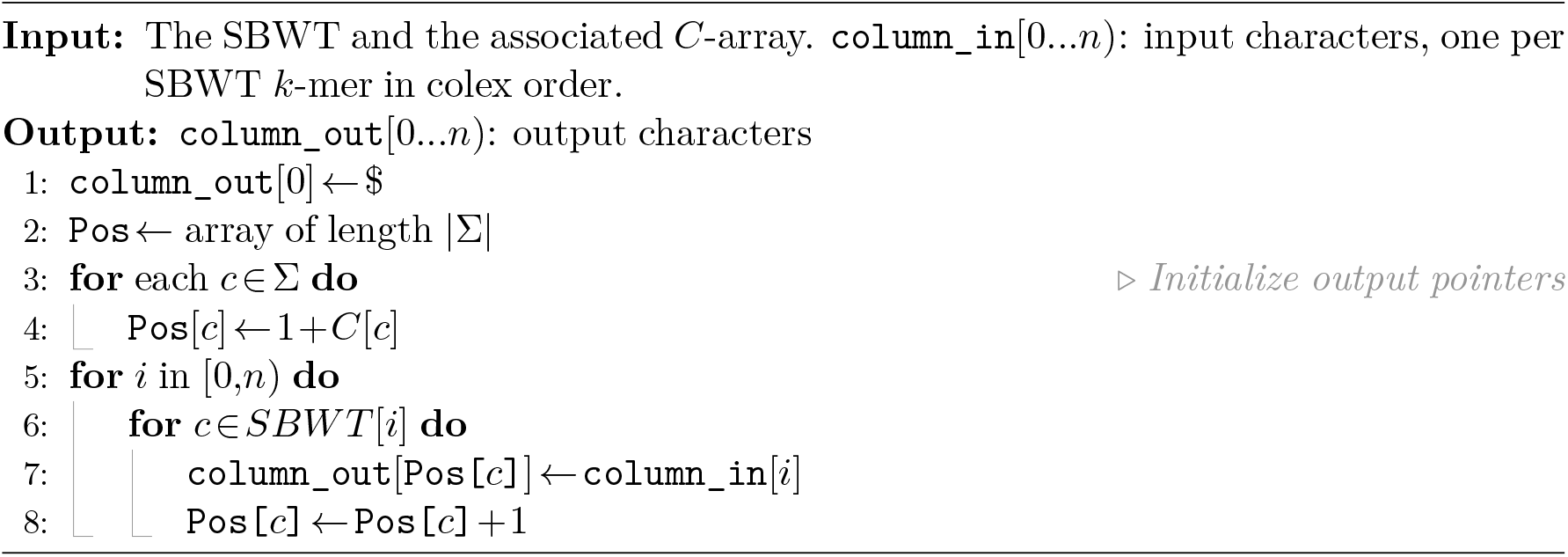

We implement both a low-memory option that encodes the labels `column_in` and `column_out` in ⌈log_2_(|∑| +1)⌉ bits per character (+1 for the special dollar symbol), and a faster high-memory option that uses the smallest possible native machine integer types to encode characters, using 8, 16, 32 or 64 bits per character.

#### 3.1.2 Computing the merge plan in *O*(*n*log*k*) time and *O*(*n*log*n*) bits

We now describe an asymptotically faster approach for computing the merge plan. We do not implement this algorithm in this paper, but include it anyway for completeness.

The algorithm works by applying the technique of prefix doubling to the SBWT graph. We denote the nodes and edges of the first input SBWT graph by (*V*_1_,*E*_1_), and those of the second input SBWT graph by (*V*_2_,*E*_2_). Both graphs are preprocessed so that the source node has a self-loop with the label $ that is smaller than all other characters. For simplicity, we assume that *k* = 2^*b*^ for some positive integer *b*, so the algorithm runs for *b* rounds (removing this assumption is trivial in practice). At round *r*, we have a mapping *π*_*r*_ : *V*_1_ ∪ *V*_2_ → ℤ sorting the nodes according to the incoming 2^*r*^-mers, that is, *π*_*r*_(*u*) =π_*r*_(*v*) if the incoming 2^*r*^-mers to *u* and *v* are the same, and otherwise π_*r*_(*u*) <π_*r*_(*v*) if the incoming 2^*r*^-mer to *u* is colexicographically smaller than the incoming 2^*r*^-mer to *v*. After the last round, the merge plan can be read from π_*b*_.

We initialize π_0_ so that π_0_(*v*) is the incoming character to *v*. Now, suppose we have constructed π_*r*_ and want to construct π_*r*+1_. Since every node in both graphs has in-degree exactly 1, it is always possible to walk backward 2^*r*^ edges unambiguously. Let back(*v*,2^*r*^) denote the node reached by walking 2^*r*^ edges backward from *v*. We assign each node *v* to the pair (π_*r*_(back(*v*,2^*r*^)),π_*r*_(*v*)). These pairs are compared by using the second element as the primary key, and the first element as the secondary key. With this comparison, the order of the pairs matches the colexicographic order of the incoming 2^*r*+1^-mers. The mapping π_*r*+1_ is obtained by radix-sorting the pairs (right-to-left), and replacing the pairs with single integers preserving the order of the pairs.

To ensure that each round takes linear time, we do not actually walk backward 2^*r*^ steps from each node, but instead we maintain pointers from *v* to back(*v*,2^*r*^) at round *r*. The back-pointers for the next round can be constructed by traversing two back-pointers from the current round.

### 3.2 From merge plans to functional SBWT indexes

Given the merge plan, constructing the bit matrix SBWT representation resulting from a set operation on two padded *k*-spectra *A*^+^ and *B*^+^ is conceptually straightforward. For a union, we keep all positions marked L, R, or B in the merge plan: rows marked L or R are copied from the corresponding input, while rows marked B are obtained by taking the bitwise OR of the two corresponding input rows. For an intersection, we keep only the positions marked B and take the bitwise AND of the two corresponding input rows. For a set difference *A*^+^\*B*^+^, we keep only the positions marked L.

The difficulty is that the SBWT does not encode only the real *k*-mers: it also contains the dummy *k*-mers introduced in the padded *k*-spectra. These dummies represent the minimal set of *k*-mers that must be introduced to guarantee that every non-dummy *k*-mer has an incoming path of length at least *k* in the SBWT graph, starting at the source node representing *k*-mer $^*k*^. When two SBWTs are combined, some correction is required to keep the dummy *k*-mer set both functional and minimal.

#### 3.2.1 Pruning redundant dummy nodes during the union operation

For the union operation, the dummy problem can be described as follows. Let 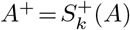 and 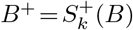 denote the padded k-spectra of the two input *k*-mer sets. A naive merge based on the merge plan yields the colexicographically sorted set *A*^+^ ∪ *B*^+^. However, the desired output is not this set, but rather the padded *k*-spectrum of the union, 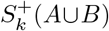. In the latter, the dummy *k*-mer set is minimal, while the former may contain redundant dummy *k*-mers. When the dummy *k*-mer set is not minimal, the SBWT graph requirement that each node has exactly one incoming-character edge is violated.

**Example**. Consider the case *k* = 5 and the two singleton sets *A*={ACTGA} and *B* ={CTGAC}. Their padded *k*-spectra are 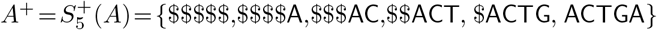 and 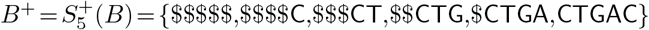. As shown in Figure 1, the naïve merge *A*^+^ ∪ *B*^+^ is not the canonical padded spectrum of the union *A* ∪ *B*. In the combined set, ACTGA already provides a predecessor for CTGAC, so dummy $CTGA is redundant. The redundancy then propagates backwards along the dummy chain, rendering $$CTG, $$$CT, and $$$$C redundant as well. Thus, 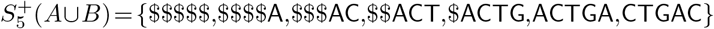.

**Figure 1.**
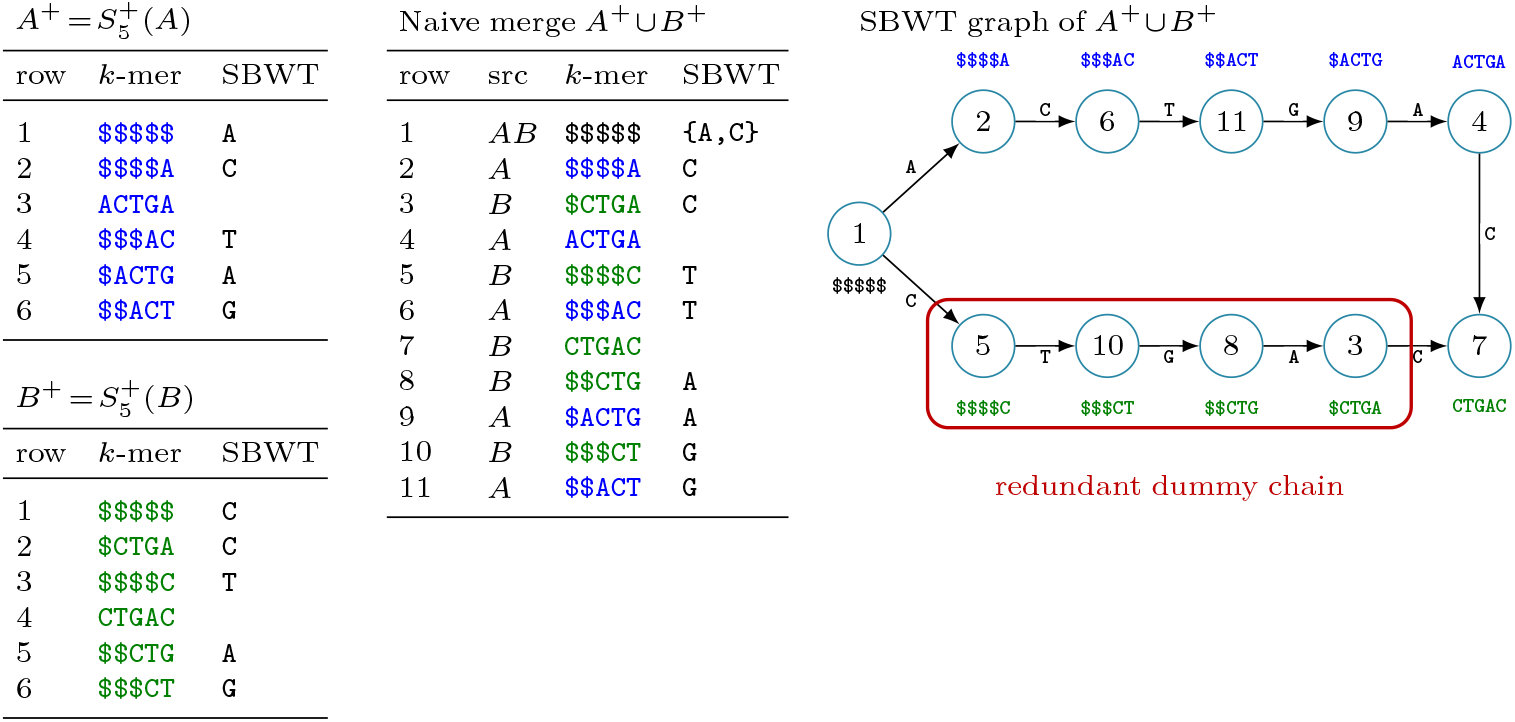
Naive SBWT merge can create redundant dummy nodes. Example with *k* = 5, *A* = {ACTGA}, and *B* = {CTGAC}. Left: the two padded spectra in colexicographic order, with corresponding SBWT subset sequence. Middle: the naive merged order and subset sequence based on the merge plan. SBWT columns, equivalently the outgoing edge labels in the SBWT graph, are encoded in an SBWT bit matrix; the *k*-mer strings are shown for clarity but not stored. Right: the SBWT graph induced by *A*^+^ ∪ *B*^+^. The lower dummy chain is redundant because ACTGA already provides a predecessor for CTGAC, so $CTGA and then recursively its dummy predecessors can be pruned.

#### Pruning approach

Having obtained the merge plan for *A*^+^ and *B*^+^ and the SBWT bit matrix corresponding to the naive union *A*^+^ ∪ *B*^+^, redundant dummy nodes can be pruned in three stages (accompanying pseudocode is shown in Algorithm S1).

*Stage 1: mark directly redundant dummies*. We scan each suffix group in *A*^+^ ∪ *B*^+^, i.e., each maximal interval of consecutive rows whose *k*-mers share the same suffix of length *k* − 1.^1^ Within a group, a dummy *k*-mer originating exclusively from *A*^+^ is *directly redundant* if the same group contains at least one real *k*-mer that is exclusive to *B*^+^, or vice versa. For example, in Figure 1, rows 3 and 4 in the merged order form one suffix group, with dummy row 3 exclusive to *B*^+^ and real row 4 exclusive to *A*^+^. In this case, the real *k*-mer already provides the predecessor that the dummy *k*-mer was introduced to supply. Let *R* be the set of redundant rows identified in this stage.

*Stage 2: propagate redundancy backwards*. Predecessor nodes in the dummy chains preceding the *k*-mers corresponding to rows in *R* may also become redundant. We therefore propagate redundancy backwards through the SBWT graph. A dummy row *p*= pred(*q*), for some *q* ∈ *R*, is added to *R* if *p* ∉ *R* and succ(*p,c*) ∈ *R* for every *c* ∈ SBWT[*p*]. That is, all successors of *p* are already redundant, so *p* carries no useful information. For example, in Figure 1, once row 3 is in *R*, its predecessor row 8 is added because its only successor is row 3, and this then propagates to rows 10 and 5. The propagation is implemented as a level-synchronous breadth-first search and terminates when a full level adds no new rows.

*Stage 3: filter the rows*. Finally, all rows in *R* are deleted from the merged SBWT bit matrix representing *A*^+^ ∪ *B*^+^. While scanning the rows, we process one suffix group at a time. Outgoing edge bits from deleted rows are accumulated, except for edges whose successor row is also in *R*. If the suffix group has a surviving row, the accumulated bits are added to the first surviving row of the group. This preserves the SBWT convention that outgoing labels are stored at the first row of each suffix group. The resulting subset sequence is the SBWT of 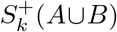.

#### 3.2.2 Adding missing dummy nodes during the intersection operation

Whereas for union the naive merge of *A*^+^ and *B*^+^ could contain redundant dummy *k*-mers, for intersection the opposite problem occurs: a naive row-wise intersection may miss dummy *k*-mers that are required for a valid SBWT index. In such a case, the character counts in the SBWT subset sequence no longer match the character counts in the last column of the colexicographically sorted matrix. Equivalently, the SBWT graph contains source nodes other than the global source $^*k*^. These extra sources are precisely the rows whose missing dummy predecessors must be added. In other words, intersection cannot be implemented by simply keeping the shared outgoing edges of suffix groups that contain *k*-mers occurring in both input padded spectra. Rather, after identifying shared *k*-mers, we must also add any dummy chains needed to make the resulting set prefix-complete.

**Example**. Consider the case *k* = 5 and the two *k*-mer sets *A* = {ACTGA,CTGAC} (from string ACTGAC) and *B* = {TCTGA,CTGAC} (from string TCTGAC). Their padded spectra are 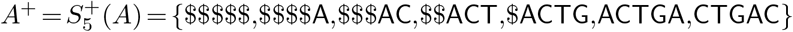 and 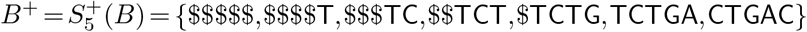. As shown in Figure 2, the naive intersection keeps only the shared *k*-mers $$$$$ and CTGAC, with no shared outgoing labels. This is not a valid SBWT index, because the dummy predecessors $CTGA, $$CTG, $$$CT, and $$$$C are missing. The canonical padded spectrum requires adding the dummy chain ending at CTGAC. Thus, 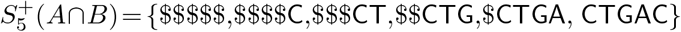.

**Figure 2.**
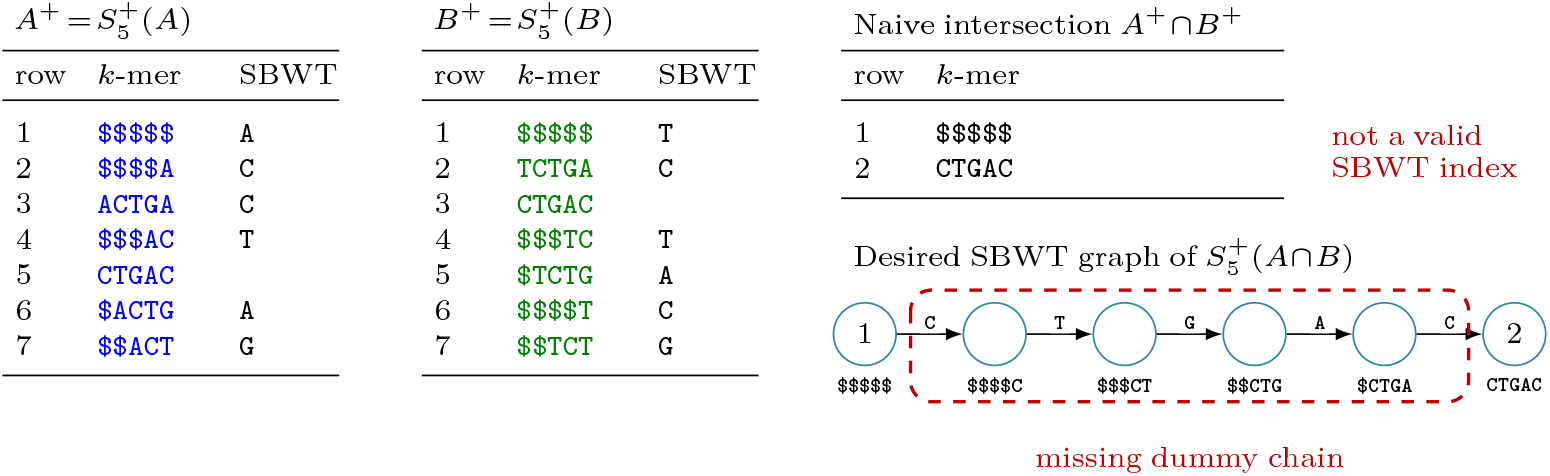
Intersection can miss required dummy rows. Example with *k* = 5, *A* = {ACTGA, CTGAC}, and *B* = {TCTGA,CTGAC}. Left: the two padded spectra in colexicographic order, with corresponding SBWT subset sequence. Top right: the naive intersection *A*^+^ ∩ *B*^+^, which keeps only $^5^ and *k*-mer CTGAC without any shared outgoing labels, and is not a valid SBWT index. Bottom right: the desired SBWT graph for 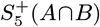, where the missing dummy chain ending at CTGAC has been added.

#### Missing dummy repair

Having obtained the merge plan for *A*^+^ and *B*^+^, we can construct 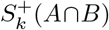 in two major stages. The first stage tests whether the naive intersection is already prefix-complete, where the naive intersection keeps only outgoing labels that appear in both inputs for suffix groups containing rows shared by both *A*^+^ and *B*^+^. If so, no repair is required. Otherwise, a dummy-repair stage reconstructs the missing predecessor chains by building a new three-way merge plan. Accompanying pseudocode is shown in Algorithm S2.

*Stage 1: incoming-edge coverage*. We scan the suffix groups of the merged order. For each suffix group that contains at least one shared *k*-mer, the naive intersection keeps exactly the outgoing labels that are present in both input SBWTs for that suffix group. For each resulting label *c*, the successor reached by following the corresponding edge in the SBWT graph is marked as having an incoming edge in the intersection. Since the naive intersection may not be a functional SBWT, the successor is computed using rank support in the *A*^+^ colexicographic space. After the scan, any non-dummy intersection *k*-mer that was not marked is a *source node* violating SBWT requirements. If no such new source nodes exist, all non-source intersection rows already have incoming edges and SBWT index construction is trivial. Otherwise, dummy-chain repair is required.

*Stage 2a: repair missing dummy chains*. Each new source node is a non-dummy *k*-mer in the intersection that lacks a predecessor. Let 𝒫 contain the colexicographic positions of all new source nodes in the *A*^+^ colexicographic space. For each position *p* ∈ 𝒫, we reconstruct the corresponding *k*-mer by starting at *p* and repeating *p* ← pred(*p*) for *k* steps in *A*^+^, recording the incoming edge label at each step. The set *P* of reconstructed source *k*-mers is then passed to the SBWT builder. This produces a third index representing the padded *k*-spectrum 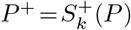, whose dummy chains supply the missing predecessors.

*Stage 2b: build the repaired intersection*. We compute a three-way merge plan of *A*^+^, *B*^+^, and *P*^+^ in colexicographic order using the *O*(*nk*) time implementation. A row contributes to the final intersection if it represents either a shared *k*-mer from *A*^+^ ∩ *B*^+^ or a row from *P*^+^. For each contributing suffix group, outgoing labels are selected by taking those present in *P*^+^ or present in both *A*^+^ and *B*^+^. The resulting subset sequence is the SBWT of 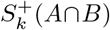.

#### 3.2.3 Adding missing dummy nodes during the set difference operation

To compute the set difference *A\B* using the merge plan, we keep rows and corresponding outgoing labels that are present in *A*^+^ but not in *B*^+^. Constructing the SBWT index in this way needs the same dummy-repair stage as for intersection. In the resulting three-way merge plan, output rows and their labels are selected by taking those present in the padded spectrum *P*^+^ or those present in *A*^+^ but not in *B*^+^. This approach differs from intersection in one additional detail. The global source $^*k*^ is present in both inputs and is thus not selected. It must therefore be inserted explicitly into the naive difference order, with outgoing labels selected as those present in *A*^+^ but not in *B*^+^ for the global source suffix group.

## 4 Experiments

This section presents experimental results demonstrating the performance of set operations on SBWT indexes. The evaluation has two parts. First, we benchmark SBWT-based union, intersection, and set difference on datasets with different overlap characteristics, reporting runtime, memory usage, and index size alongside two baseline methods: KMC [13] and a straightforward in-house sorted-list implementation. Second, we show that SBWT merging can serve as a practical building block for large-scale index construction by building an SBWT index for the complete 661K bacterial genomes dataset [6] using a binomial merge pipeline.

### 4.1 Benchmarking SBWT set operations

#### Data

We benchmarked set operations on two pairs of datasets chosen to represent different overlap regimes. For a high-overlap setting, we used paternal assembly GCA_018472715.2 and maternal assembly GCA_018472765.3. These correspond to sample HG00735 from the Human Pangenome Reference Consortium [19]. Because human genomes are highly conserved at the sequence level, the *k*-mer spectra of the two haplotype assemblies are expected to overlap strongly. These assemblies were indexed on the forward strand only.

For a lower-overlap setting, we used two diverse sequence collections: the first batch of 1 000 assemblies from the ENA 661K bacterial genomes dataset [6], and the complete PLSDB plasmid database [18]. Because these datasets combine sequences from heterogeneous public sources, both strands were indexed to avoid bias due to assembly or plasmid orientation. Additional dataset statistics are detailed in Supplementary Table S1.

#### Environment

All benchmarks in this subsection were run on a server with an Intel Xeon Gold 6430 CPU, 64 cores, and 503.4 GiB of RAM. Each configuration was run once as a warm-up and then repeated 10 times; reported runtimes are medians over the 10 repetitions.

#### Baselines

We compared SBWT set operations^2^ in both default and low-memory modes (Section 3.1.1) to KMC version 3.2.4 and a straightforward in-house sorted-list baseline^3^. All tools were run with *k* = 31. We do not include dynamic *k*-mer set structures [14, 1] in our experiments because they have a different design goal than us (i.e., support for insertion and deletion of individual *k*-mers) and are slow at bulk operations.

KMC is a *k*-mer counting tool that stores distinct *k*-mers, optionally with counts, in a compact database representation. KMC3 builds its database in two stages: first, input *k*-mers are partitioned into bins using short signatures; second, the bins are sorted independently to remove duplicates and compute counts [13]. The set operations then follow a merge-like scan over ordered *k*-mer streams from the two input databases.

The sorted-list baseline encodes each *k*-mer as a 64-bit integer using 2 bits per base, which supports *k* ≤ 32. During construction, *k*-mers (and, when requested, their reverse complements) are extracted from the input sequences, sorted with a radix sort, and deduplicated, yielding a sorted array of distinct *k*-mers. Input parsing is performed using the Helicase library [15].

The resulting sets are stored in a compact binary format. Set operations are implemented by a standard two-pointer merge scan over the two sorted arrays, conceptually similar to the approach used by KMC. The scan is performed through buffered iterators over the sorted arrays on disk, so the inputs need not be fully loaded into memory.

We emphasize that KMC and the sorted-list implementation serve as *baselines* for exact *k*-mer set operations. Both represent sets in forms that are well suited to ordered scans and database-style set manipulation. The SBWT has a different goal: it represents the set as a compressed index that supports membership queries in time independent of the number of indexed *k*-mers, and navigation between overlapping *k*-mers in the SBWT graph. Thus, the comparison below is aimed at measuring the overhead of preserving this richer indexed structure while performing set operations, rather than comparing interchangeable structures.

#### 4.1.1 High-overlap benchmark: human haplotypes

Table 1 reports the build and set-operation statistics for the human haplotype benchmark. The set-operation measurements from the table (together with supplementary thread counts) are visualized in Figure 3, which shows the runtime-memory trade-off for union, intersection, and set difference. As reported in Table 1, the two assemblies contain almost the same number of distinct 31-mers. As expected in this high-overlap setting, the intersection is large, the difference is much smaller, and the union remains close in size to the two input sets.

**Table 1.** Performance metrics for building and performing set operations on 31-mer indexes of two human haplotypes (HG00735). Values represent the median of 10 repetitions. SBWT and KMC used 64 threads; the sorted-lists baseline was single-threaded. SBWT is slower than the baselines for set operations, but produces compressed indexes that are about an order of magnitude smaller than those of the baseline tools, while preserving a queryable and navigable representation of the *k*-mer set. Detailed command-line settings are given in Supplementary Table S2.

|  | Assemblies |  | Set Operations |  |  |
| --- | --- | --- | --- | --- | --- |
| | paternal | maternal | pat. ∪ mat. | pat. ∩ mat. | pat. $\setminus$ mat. |
| # 31-mers (Millions) | 2549.46 | 2548.23 | 2635.99 | 2461.70 | 87.76 |
| **SBWT ($t_{64}$)** |  |  |  |  |  |
| Index size (GiB) | 1.48 | 1.48 | 1.54 | 1.46 | 0.08 |
| Runtime (s) | 59.49 | 60.19 | 94.66 | 188.59 | 190.49 |
| low-memory |  |  | 148.74 | 295.66 | 294.80 |
| Peak RAM (GiB) | 27.65 | 27.66 | 16.78 | 21.04 | 21.12 |
| low-memory |  |  | 9.84 | 13.73 | 14.16 |
| **KMC ($t_{64}$)** |  |  |  |  |  |
| Index size (GiB) | 19.00 | 18.99 | 14.76 | 13.79 | 0.52 |
| Runtime (s) | 36.96 | 36.70 | 26.99 | 26.69 | 18.75 |
| Peak RAM (GiB) | 30.50 | 30.33 | 7.27 | 7.28 | 7.28 |
| **Sorted lists ($t_1$)** |  |  |  |  |  |
| Index size (GiB) | 18.99 | 18.99 | 19.64 | 18.34 | 0.65 |
| Runtime (s) | 194.93 | 196.35 | 58.99 | 58.37 | 25.04 |
| Peak RAM (GiB) | 45.48 | 45.56 | 19.64 | 18.34 | 0.66 |

**Figure 3.**
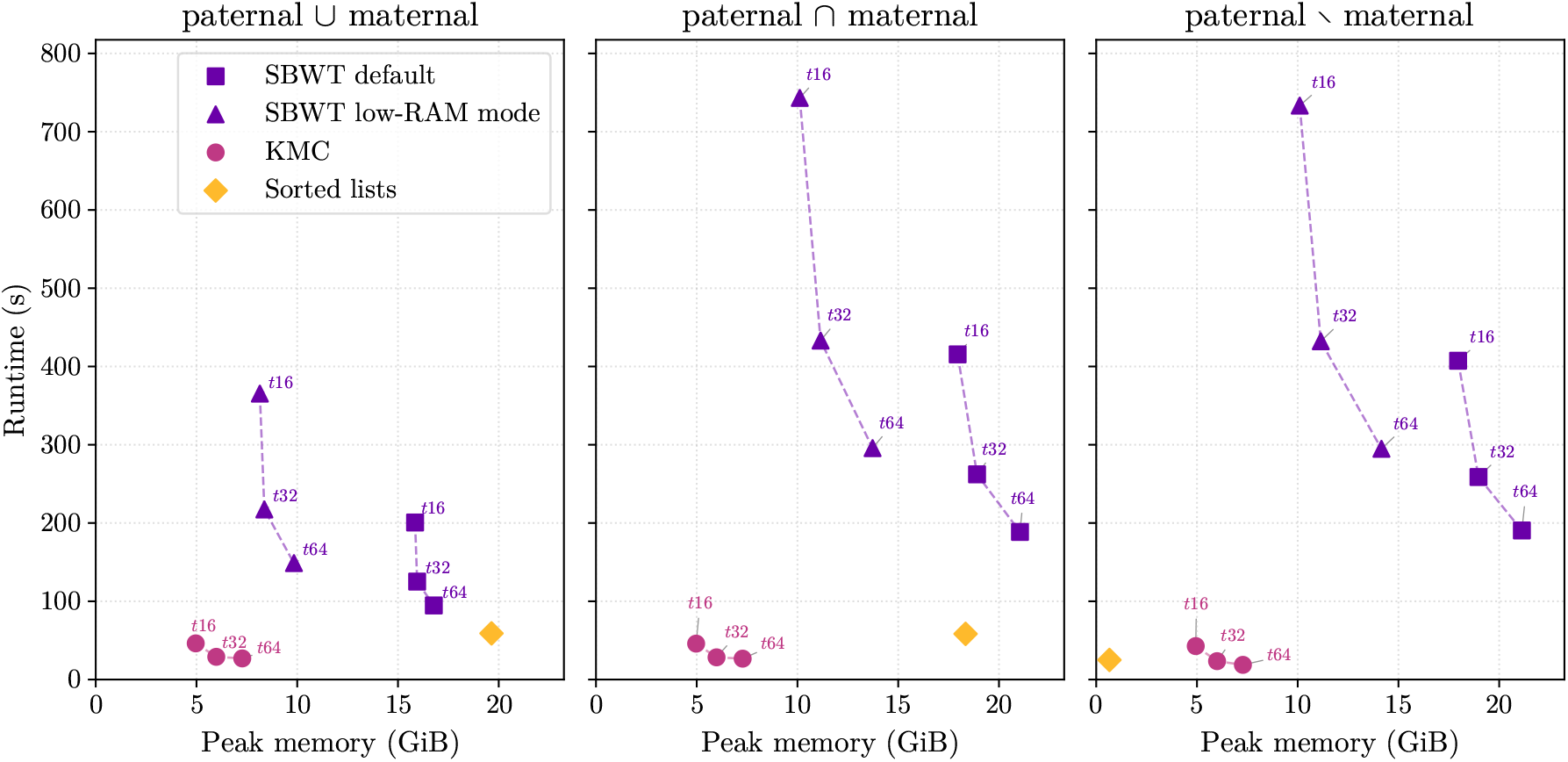
Runtime vs. peak memory for three 31-mer set operations on the paternal and maternal haplotype assemblies of sample HG00735. Thread counts of 16, 32, and 64 are shown for the multi-threaded tools (KMC, SBWT default, and SBWT low-memory mode). The sorted-lists baseline is single-threaded. The *t*64 values correspond to the set-operation columns of Table 1. The baseline tools (KMC and sorted lists) perform ordered *k*-mer set manipulation, whereas SBWT set operations additionally construct a richer indexed representation supporting *k*-mer lookup and SBWT graph navigation; this extra indexing work explains the higher runtime and memory usage.

The most striking difference between the three methods is index size. For both input haplotypes, the SBWT indexes are about an order of magnitude smaller than the corresponding KMC and sorted-list representations. The same pattern holds approximately for the set-operation outputs. This reflects one main benefit of the SBWT representation: the output is a *compressed* index of the *k*-mer set, rather than an ordered *k*-mer database or list.

When comparing set-operation performance, KMC (using 64 threads) is the fastest method for all three set operations. The sorted-list baseline is somewhat slower but competitive despite being single-threaded. Relative to KMC, default SBWT is 3.5× slower for union, 7.1× slower for intersection, and 10.2× slower for difference. This overhead can be explained by the fact that the SBWT algorithms operate on compactly encoded indexes and must construct a valid SBWT index (including the dummy-node corrections described in Section 3.2), rather than merely reading and writing ordered collections of naively encoded *k*-mers. Profiling on the human benchmark shows that merge-plan construction (Section 3.1.1) dominates SBWT set-operation runtime, accounting for 86.7–93.5% of union time and 93.1–97.2% of intersection and difference time (including the three-way merge plan used for dummy repair).

KMC uses the least RAM during the set operations. The sorted-list baseline has peak memory usage close to its output size, because the scan itself is streaming but the output set is stored explicitly in memory. This explains why sorted lists use little memory for the small-difference output, but substantially more memory for the much larger union and intersection outputs. For SBWT, intersection and difference use more memory than union because these operations require a three-way merge plan construction, whereas union uses only the two-way merge plan. The low-memory SBWT mode (see Section 3.1.1) reduces peak memory significantly relative to the default mode. This memory reduction comes at the cost of increased runtime.

#### 4.1.2 Low-overlap benchmark: bacteria and plasmids

Table 2 and Figure 4 show statistics for the bacteria–plasmid benchmark. In contrast to the human haplotypes, these datasets have little overlap: the intersection is small, the union is close to the sum of the two inputs, and most bacterial 31-mers are absent from the plasmids.

**Table 2.**
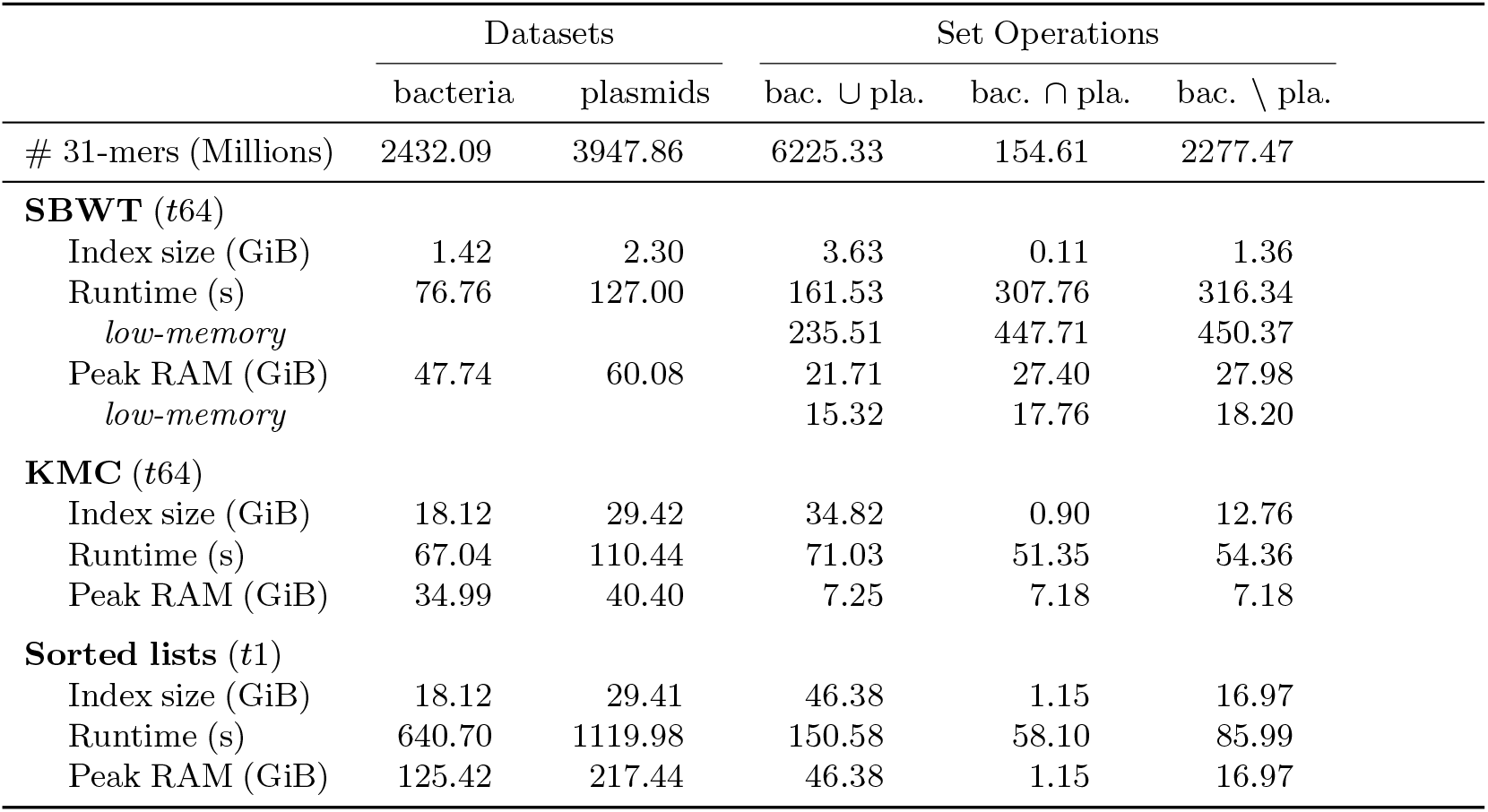
Performance metrics for building and performing set operations on 31-mer indexes for 1 000 bacterial assemblies and a plasmid dataset. Values represent the median of 10 repetitions. SBWT and KMC used 64 threads; the sorted-lists baseline was single-threaded. SBWT is slower than the baselines for set operations, but produces compressed indexes that are about an order of magnitude smaller than those of the baseline tools, while preserving a queryable and navigable representation of the *k*-mer set. Detailed command-line settings are given in Supplementary Table S2.

|  | Datasets |  | Set Operations |  |  |
| --- | --- | --- | --- | --- | --- |
| | bacteria | plasmids | bac. ∪ pla. | bac. ∩ pla. | bac. $\setminus$ pla. |
| # 31-mers (Millions) | 2432.09 | 3947.86 | 6225.33 | 154.61 | 2277.47 |
| **SBWT ($t_{64}$)** |  |  |  |  |  |
| Index size (GiB) | 1.42 | 2.30 | 3.63 | 0.11 | 1.36 |
| Runtime (s) | 76.76 | 127.00 | 161.53 | 307.76 | 316.34 |
| *low-memory* |  |  | 235.51 | 447.71 | 450.37 |
| Peak RAM (GiB) | 47.74 | 60.08 | 21.71 | 27.40 | 27.98 |
| *low-memory* |  |  | 15.32 | 17.76 | 18.20 |
| **KMC ($t_{64}$)** |  |  |  |  |  |
| Index size (GiB) | 18.12 | 29.42 | 34.82 | 0.90 | 12.76 |
| Runtime (s) | 67.04 | 110.44 | 71.03 | 51.35 | 54.36 |
| Peak RAM (GiB) | 34.99 | 40.40 | 7.25 | 7.18 | 7.18 |
| **Sorted lists ($t_1$)** |  |  |  |  |  |
| Index size (GiB) | 18.12 | 29.41 | 46.38 | 1.15 | 16.97 |
| Runtime (s) | 640.70 | 1119.98 | 150.58 | 58.10 | 85.99 |
| Peak RAM (GiB) | 125.42 | 217.44 | 46.38 | 1.15 | 16.97 |

**Figure 4.**
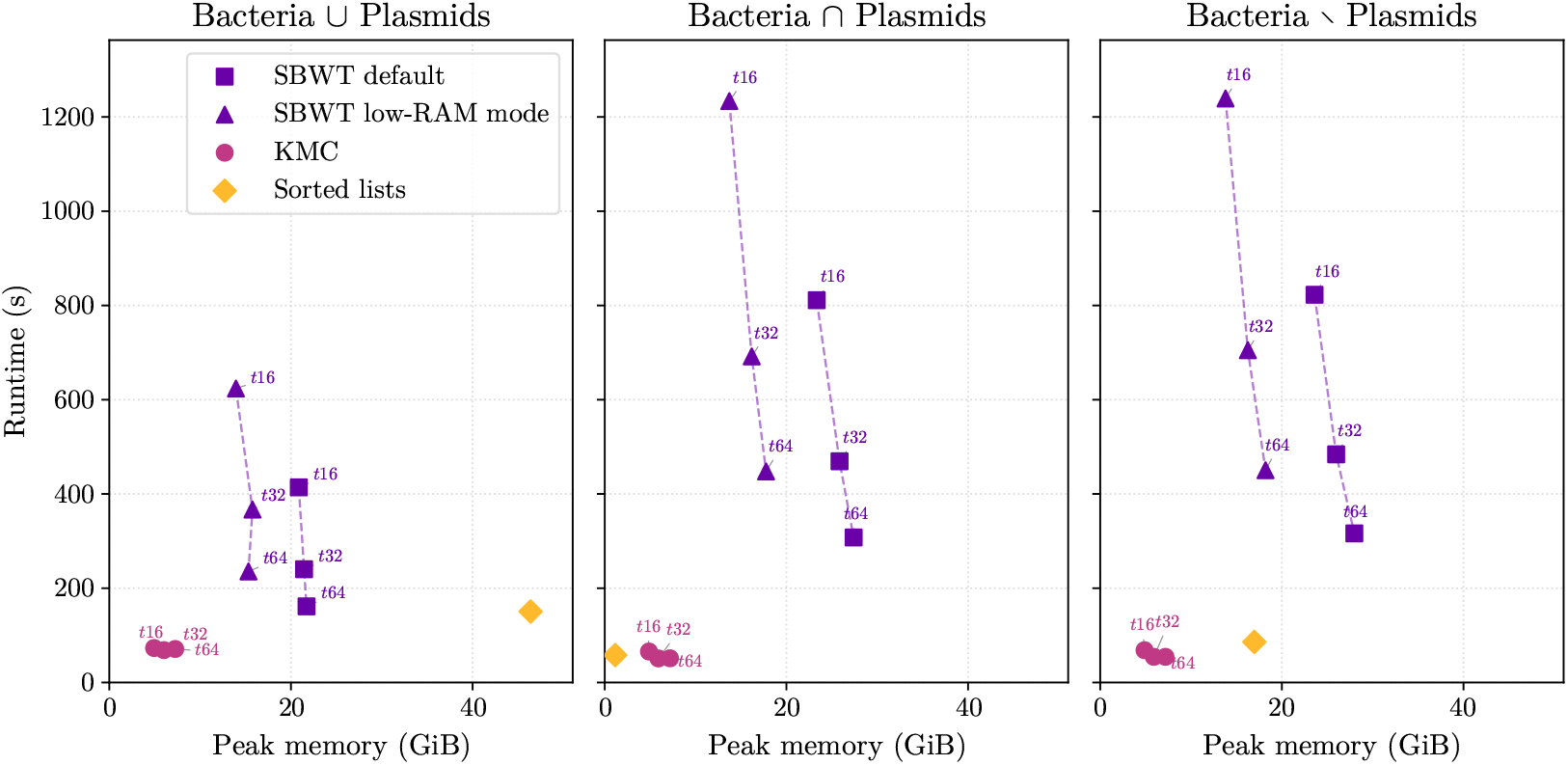
Runtime vs. peak memory for three 31-mer set operations on 1 000 bacterial assemblies and a plasmid dataset. Each point reports the median over 10 timed repetitions; thread counts of 16, 32, and 64 are shown for the multi-threaded tools (KMC, SBWT default, and SBWT low-memory mode). The sorted-lists baseline is single-threaded. The *t*64 values correspond to the set-operation columns of Table 2. The baseline tools (KMC and sorted lists) perform ordered *k*-mer set manipulation, whereas SBWT set operations additionally construct a richer indexed representation supporting *k*-mer lookup and SBWT graph navigation, which explains the higher runtime and memory usage.

The same overall pattern is observed as in the high-overlap benchmark: SBWT produces substantially smaller indexes than both baselines, KMC is again the fastest method for all three set operations, and relative to KMC, default SBWT is 2.3× slower for union, 6.0× slower for intersection, and 5.8× slower for difference. Runtime for the sorted-list baseline is close to SBWT for union, but lower for intersection and difference, where the output is smaller. As in the human benchmark, merge-plan construction dominates the SBWT runtime, accounting for 84.2–90.0% of union time and 93.9–97.2% of intersection and difference time.

The memory trends are also consistent with the human benchmark: KMC uses the least RAM during set operations, sorted lists have peak memory close to output size, and SBWT intersection and difference require more memory than union because they use a three-way merge plan construction. As before, the low-memory SBWT mode reduces peak memory relative to the default mode at the cost of increased runtime.

### 4.2 Building the 661K SBWT index through a binomial merge pipeline

#### Data

We evaluate large-scale SBWT construction through a binomial merge pipeline on the curated ENA 661K bacterial genomes dataset [6]. The full input contains 2.57 Tbp of sequence data across 661 405 assemblies. ENA distributes this dataset in 662 batches of 1 000 assemblies.

#### Environment

Most of the binomial merge pipeline was run on a server with two Intel Xeon Gold 6130 CPUs, 32 physical cores, and 128 GiB of RAM. The final merge was run on the server used for the set-operation benchmarks (Intel Xeon Gold 6430, 64 physical cores, 503.4 GiB RAM), because it required more memory. The full pipeline was run once.

#### Binomial merge pipeline design

The goal of the pipeline is to avoid constructing the full 661K index in one monolithic build, which is expensive in terms of either RAM or temporary disk usage. Instead, we gradually construct SBWT indexes for the ENA batches of 1 000 assemblies and merge these indexes incrementally. This turns large-scale construction into a sequence of smaller batch builds and pairwise merge operations.

We organize the pairwise merges using an online binomial merge stack. Each stack level stores at most one SBWT index: level 0 stores one batch, level 1 stores a merge of two batches, level 2 stores a merge of four batches, and so on. Whenever a newly built or newly merged index reaches an occupied level, the two indexes at that level are merged and carried to the next level. After all batches have been processed, the remaining stack entries are flushed by pairwise merges until a single final index remains.

In the implementation, the pipeline uses *k* = 31 and indexes both strands. For each ENA batch, the corresponding FASTA files are prefetched in the background, an SBWT index is built, and the downloaded FASTA files are deleted after construction. Batch indexes are built with 32 threads using the in-memory SBWT construction algorithm and a RAM usage target of 48 GiB. Merges are run with 32 threads. The low-memory merge mode is used for merge levels at least 5, or as a fallback if a default merge fails. See also Table S3 for command templates.

Table 3 summarizes the full 661K pipeline. The final index contains 88.70 billion distinct 31-mers and occupies 51.73 GiB on disk. The full pipeline completed in 50h 31m 13s, with batch construction accounting for 38.7% of the measured runtime and merges accounting for the remaining 61.3%. Peak memory usage occurred in the final root merge.

**Table 3.**
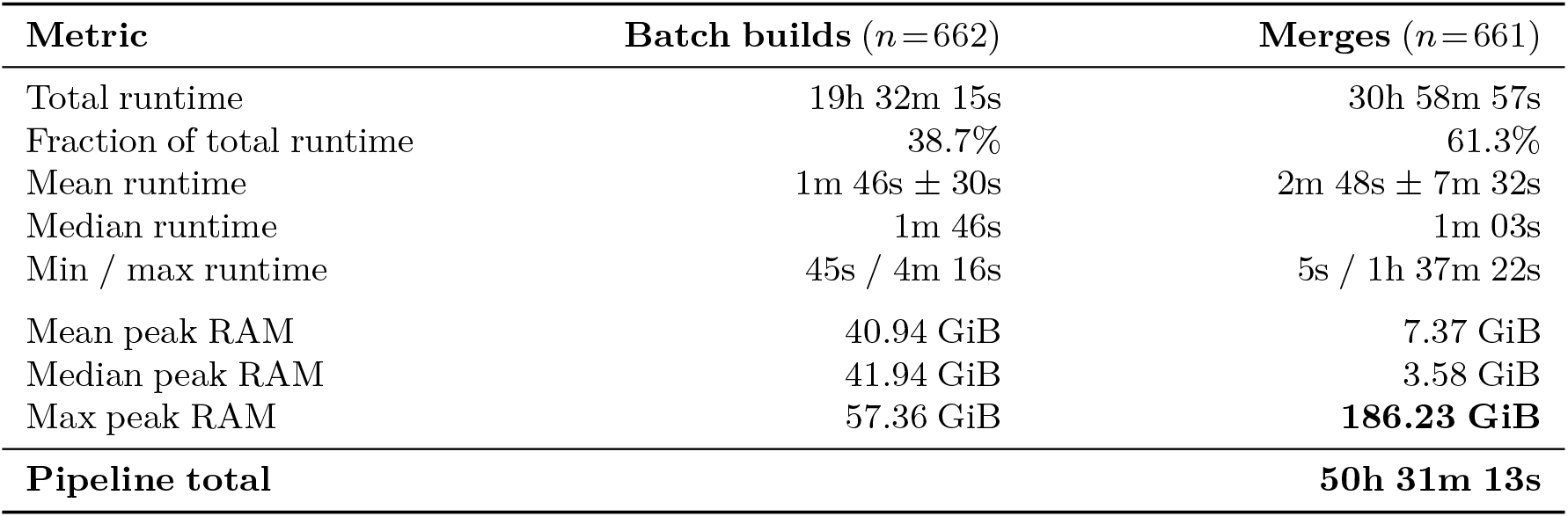
Summary of the binomial merge pipeline for construction of the SBWT index of the 661K bacterial genomes dataset (2.57 Tbp). The final root index contains 88.70 × 10^9^ distinct 31-mers and occupies 51.73 GiB on disk. The pipeline performed 662 batch builds and 661 pairwise merges.

#### 4.2.1 Comparison to direct SBWT construction

We compared the merge pipeline against direct SBWT construction [5] on prefixes of the 661K dataset. For each prefix, we used the first 2^*i*^ ENA batches, as far as computing resources allowed. We compared the binomial merge pipeline to both the in-memory SBWT construction algorithm and the disk-assisted construction algorithm. For the binomial merge pipeline, the cumulative runtime for a prefix of 2^*i*^ batches is computed as the sum of all batch-build and merge times in the corresponding merge subtree. Peak RAM is the maximum resident set size observed over all steps in that subtree. Peak temporary disk usage is the sum of index sizes of intermediate SBWT files that coexist during the binomial merge process.

Figure 5 shows the scaling behavior of the binomial merge pipeline compared with direct construction on the dataset prefixes and demonstrates why the merge pipeline is necessary at this scale when memory is limited. Direct in-memory construction has lower runtime on the prefixes where it fits in memory, but its peak RAM increases rapidly with the number of distinct 31-mers. Conversely, disk-assisted construction reduces RAM pressure, but requires substantially more temporary disk space. Both direct-construction curves stop once the corresponding resource requirement exceeds the RAM or disk-space limits on our machine.

**Figure 5.**
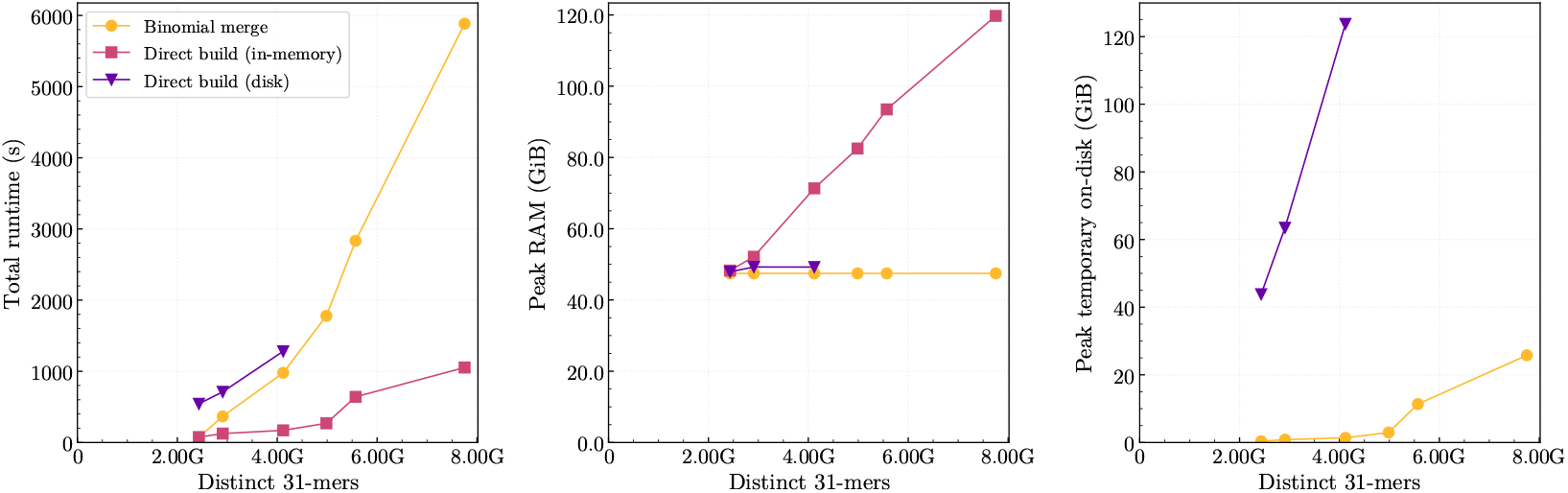
Scaling behavior of the binomial merge pipeline versus direct SBWT construction [5] as a function of the number of distinct 31-mers in the resulting index, evaluated on prefixes of the 661K dataset consisting of the first 2^*k*^ (*k* = 0,1,…,5) ENA batches of 1 000 bacterial assemblies. The direct-construction curves stop where the corresponding method exceeded the available RAM or temporary-disk limits. Left: total runtime. Centre: peak RAM usage. Right: peak temporary on-disk storage. The binomial merge pipeline is slower than direct in-memory construction on the tested prefixes, but keeps RAM and temporary-disk usage within practical limits.

The binomial merge pipeline trades some additional runtime for bounded memory and disk usage. Over the prefixes shown in Figure 5, peak RAM remains close to the per-batch construction limit, and temporary disk usage grows much more slowly than for the disk-assisted direct build. For larger indexes, merge memory eventually becomes the dominant memory cost; this transition is visible in the full merge-tree analysis in Figure 7. A rough linear extrapolation from the direct-build measurements suggests that constructing the full 661K SBWT index directly would require on the order of 1.2 TiB of RAM with the in-memory algorithm, or 5.6 TiB of temporary disk space with the disk-assisted algorithm—prohibitive amounts for most servers. By contrast, the binomial merge pipeline completed the full construction in 50h 31m 13s, with a peak RAM usage of 186.23 GiB and a peak disk usage of 112.09 GiB. This makes the binomial merge pipeline the most *practical* construction strategy for the full dataset on this hardware.

#### 4.2.2 Merge-tree scaling and resource distribution

Figure 6 shows how the number of distinct 31-mers grows across all nodes of the binomial merge tree. Each leaf corresponds to one ENA batch of 1 000 assemblies, and each internal node corresponds to a merged SBWT index. As merge depth increases, the points move to the right and upward, reflecting more input sequence and more distinct 31-mers. However, the horizontal growth is stronger than the vertical growth, showing that distinct 31-mers accumulate sublinearly with input sequence length. Thus, as more batches are merged, increasing amounts of input sequence are represented by comparatively fewer new 31-mers. At the root, the final index contains 88.70 billion distinct 31-mers over 2.57 Tbp of input sequence, corresponding to approximately one distinct 31-mer per 29 input bases.

**Figure 6.**
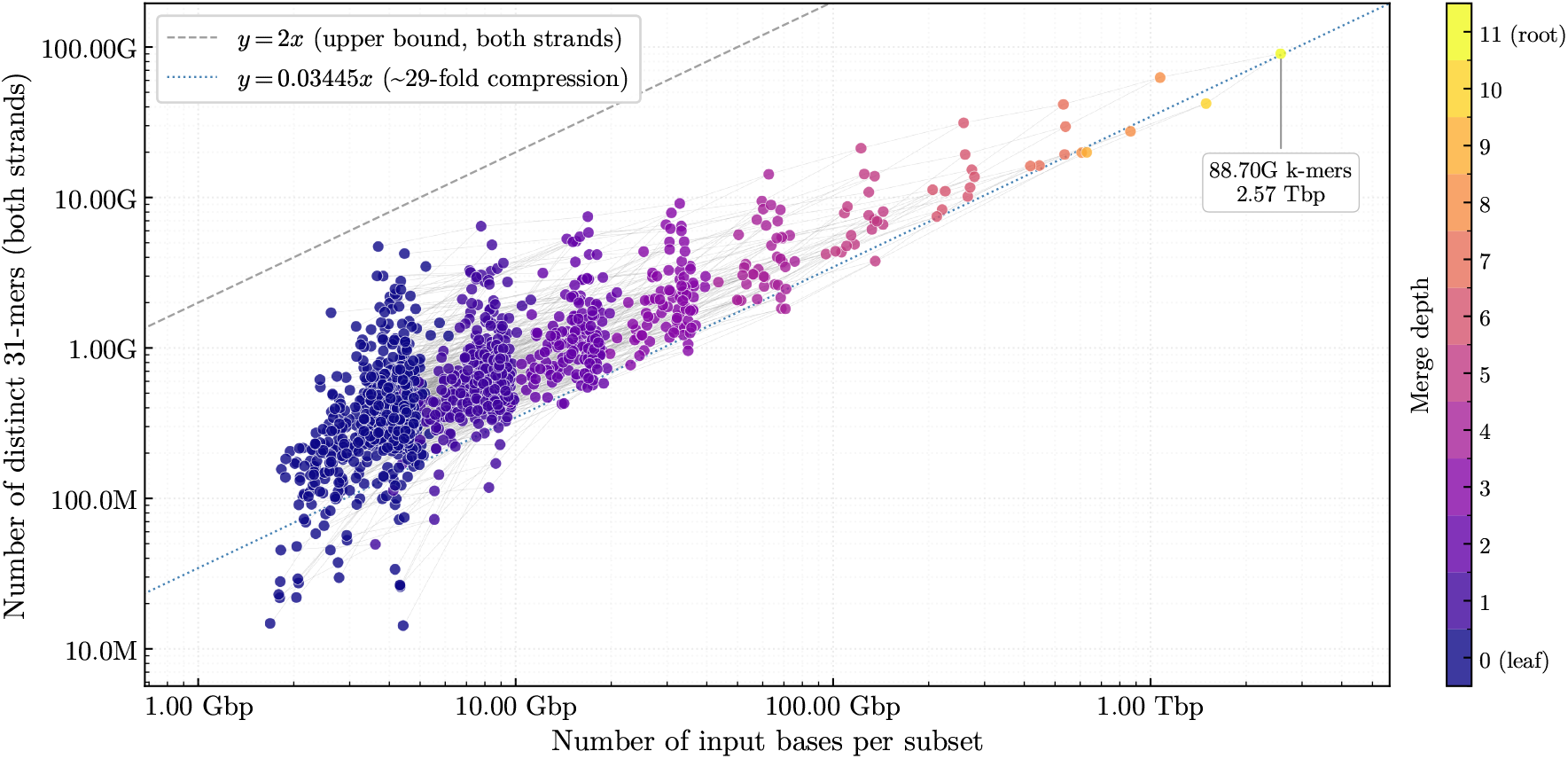
Scaling analysis of the number of distinct 31-mers as a function of total input sequence length for all nodes in the SBWT binomial merge tree for the 661K dataset. Both axes are logarithmic. Each point represents one node: either a single-batch leaf (1 000 assemblies) or a merged internal node. Color encodes merge depth in the tree. Internal nodes are connected to their two children, tracing the successive pairwise merges of the binomial tree. The dashed line *y* = 2*x* is the theoretical upper bound on the number of distinct 31-mers when indexing both strands. The dotted line *y* = 0.03445*x* shows the empirical rate at which distinct 31-mers accumulate in the full dataset: approximately one distinct 31-mer per 29 input bases. Distinct 31-mers accumulate much more slowly than input sequence length, reflecting substantial redundancy in the dataset.

Figure 7 shows where time and memory are spent across the merge tree. Batch builds have relatively stable peak memory, because they use a fixed memory target, while their runtimes vary somewhat due to batch size and sequence content. Merge operations show a clear dependence on the number of distinct 31-mers in the indexes being merged: both runtime and memory increase with merge size. Small and medium merges are inexpensive compared with batch construction, while the largest merge levels dominate the tail of the runtime and determine the peak memory requirements of the full pipeline. The low-memory merge mode appears at larger merge sizes and higher merge depths. As expected, low-memory mode shifts points toward higher runtime in exchange for lower memory usage.

**Figure 7.**
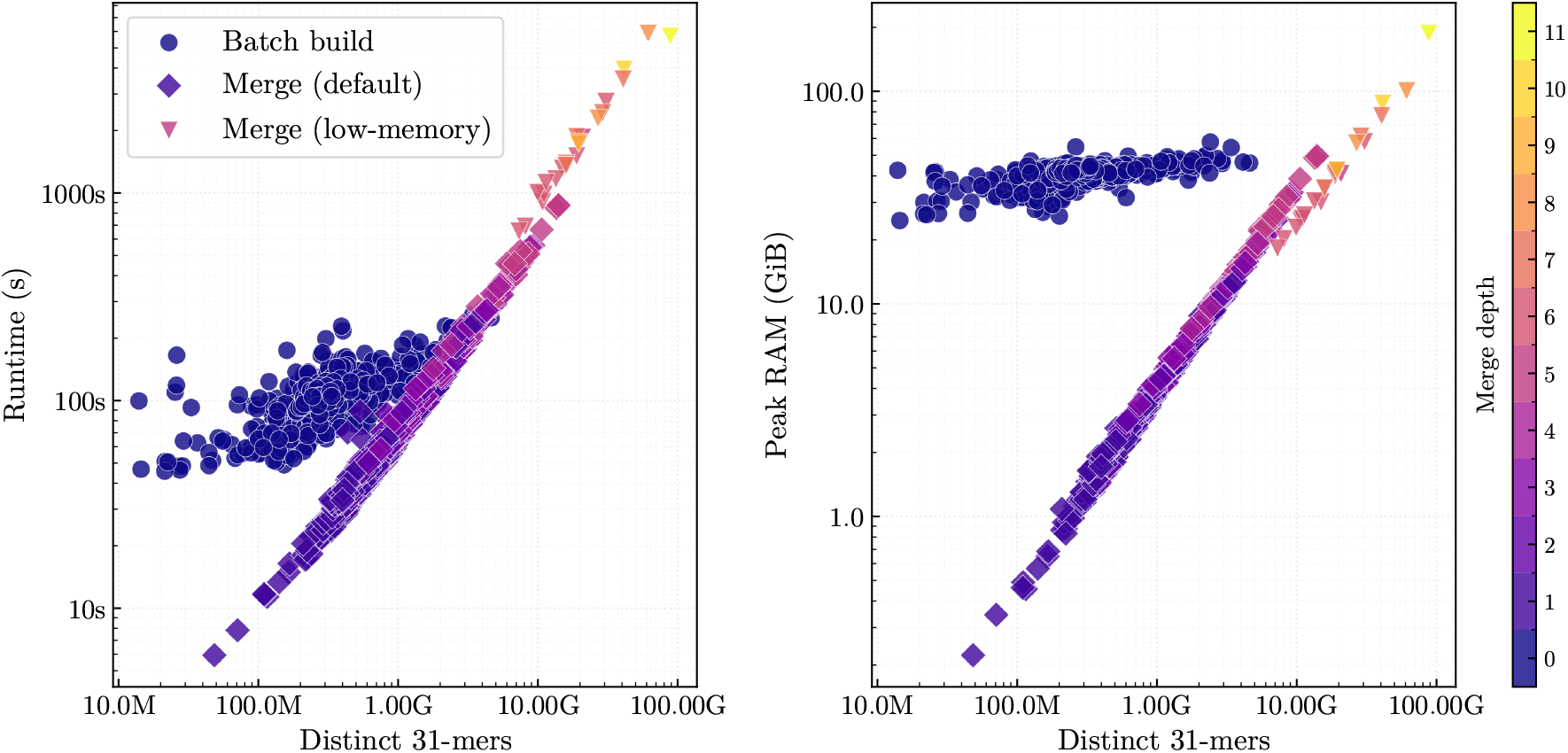
Runtime and peak memory usage as a function of the number of distinct 31-mers for all nodes in the SBWT binomial merge tree for the 661K dataset. Each point represents one pipeline stage: a single-batch index build or a pairwise merge operation. Color encodes merge depth. Left: runtime in seconds. Right: peak resident set size in GiB. Both axes are logarithmic. Batch builds have nearly fixed peak memory due to the construction memory target (48 GiB), whereas merge time and memory cost increase with *k*-mer set size. Low-memory merges are distinguished because they trade runtime for reduced memory pressure; they appear for the largest *k*-mer counts, where standard merging exceeded predefined RAM limits, or above merge level 5.

## 5 Concluding Remarks

We have presented practical algorithms for the three fundamental set operations, union, intersection, and set difference, on *k*-spectra represented as spectral Burrows-Wheeler transforms. At the core of our approach is a compact 𝒪 (*n*)-bit *merge plan*, constructible in 𝒪 (*nk*) time, together with linear-time post-processing stages that repair the dummy padding so that the output is itself a valid, queryable SBWT index. As an immediate consequence, we have described a more scalable SBWT construction algorithm.

There is much room for future work. On a technical front, our most pressing follow-up experiment is to gauge the effect of the batch size at the leaves of the merge tree on overall runtime performance. It would also be interesting to see if the *O*(*n*log*k*) algorithm we have described translates into a practical improvement in set operations, especially because merge-plan construction accounts for the largest fraction of SBWT set-operation runtime in our experiments. Also, while the 661K dataset is already of non-trivial size, in the full version of this paper we plan to include experiments with even larger datasets, such as AllTheBacteria [11]. The merge plan labels could also be repurposed for indexing *k*-mers with provenance information.

Probably the most interesting theoretical problem we leave open is the existence, or non-existence, of an *O*(*n*)-time *k*-mer set union algorithm. Recent work by Alanko and Pérez-López [4], which considers the tangentially related problem of counting *k*-mers in deterministic Wheeler graphs, suggests this may be difficult. The two counting algorithms they describe take *O*(*nk*) or *O*(*n*^4^log*k*) time. Applying these counting algorithms to a union would require determinizing the combined Wheeler graph of two *k*-mer sets, with potentially exponential blow-up; whether the SBWT avoids this is open.

## Algorithm S1

Pruning redundant dummy rows from the naively merged SBWT of *A*^+^ ∪*B*^+^

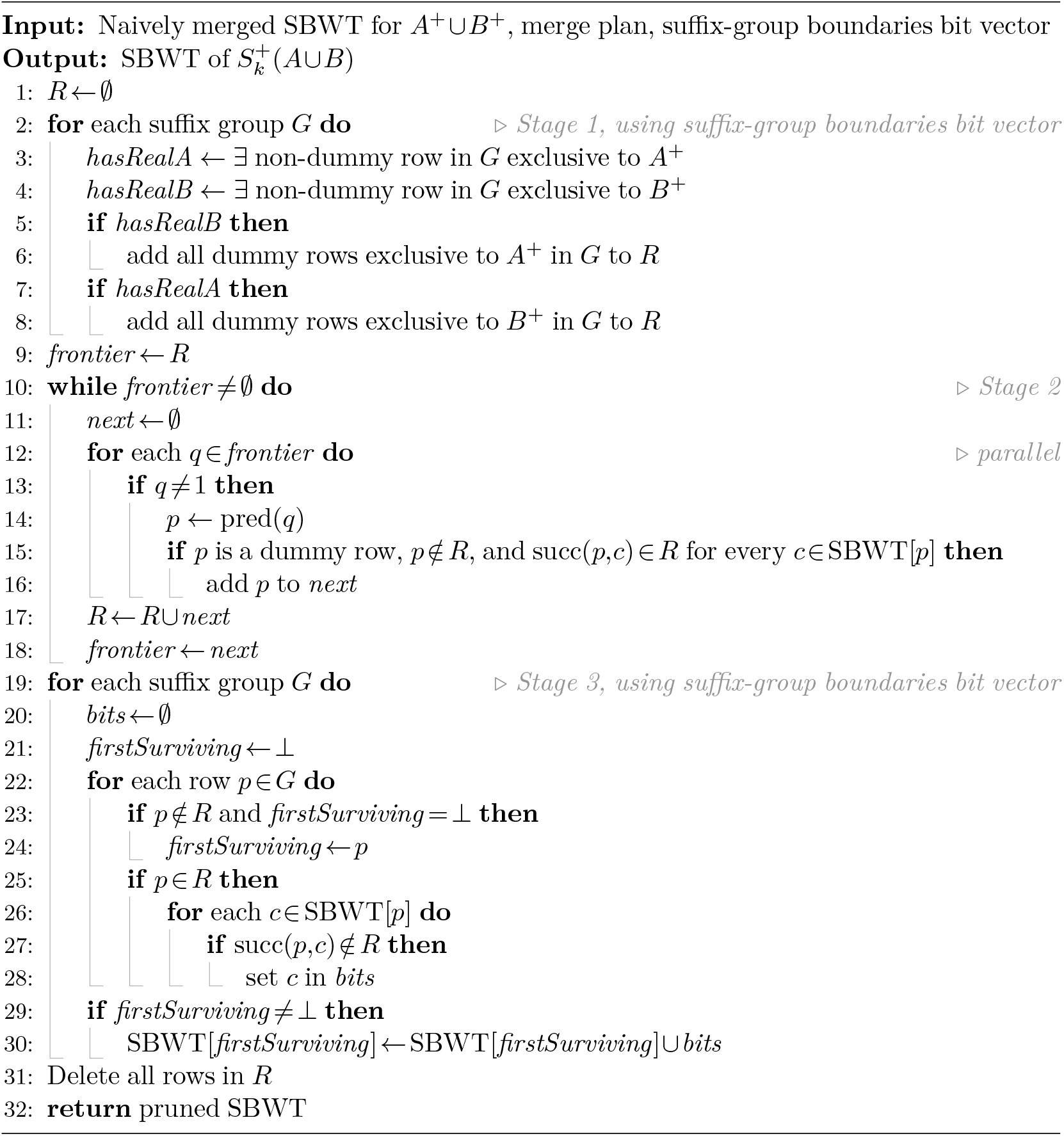

## Algorithm S2

Constructing the SBWT intersection from the merge plan of *A*^+^ and *B*^+^

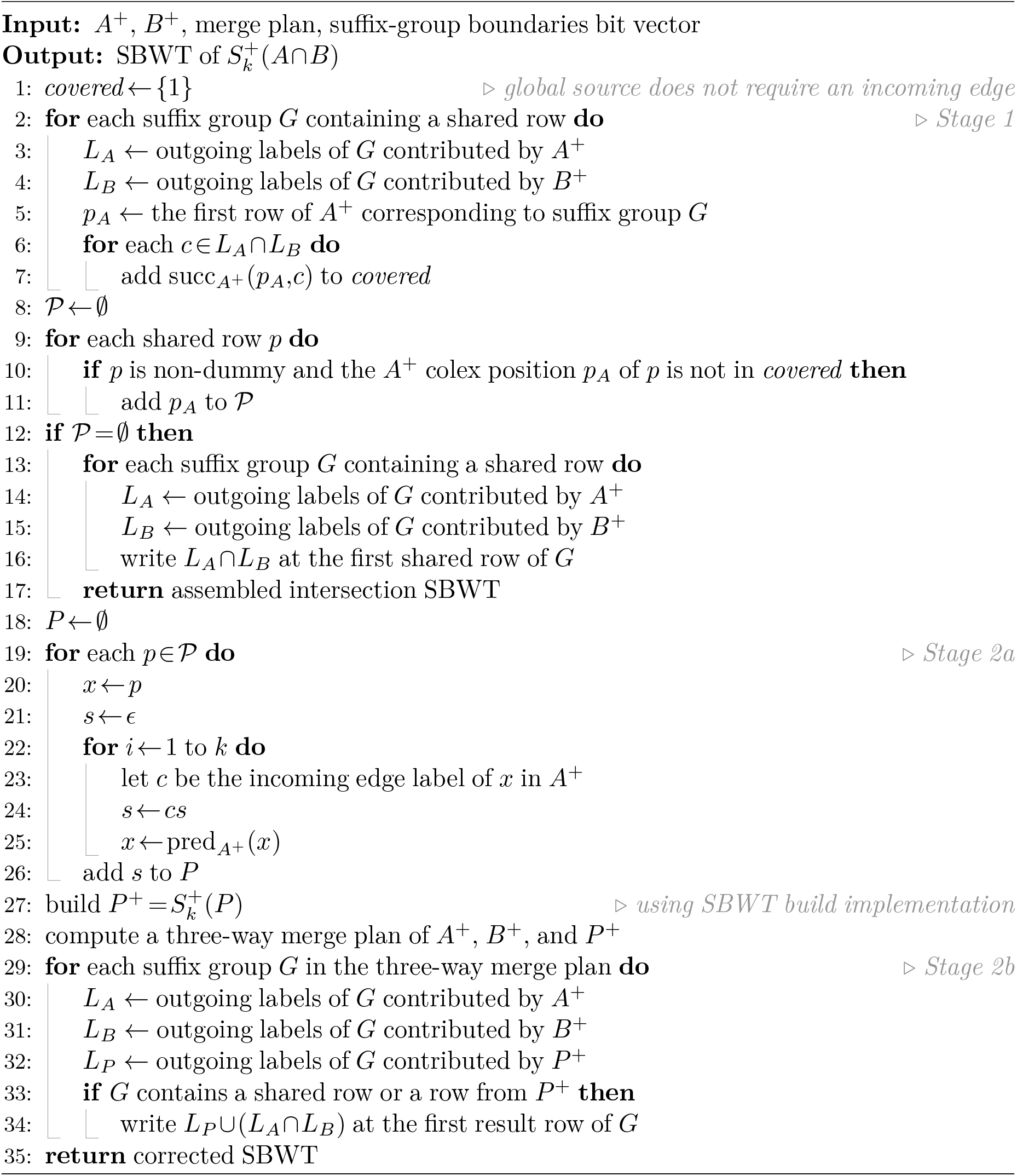

**Table S1.** Input datasets used for the set-operation benchmarks. The table reports the number of input assemblies or records, total sequence length, and gzip-compressed FASTA size.

| Dataset | Assemblies/records | Total length | Compressed size |
| --- | --- | --- | --- |
| HG00735 maternal haplotype | 1 | 3.03 Gbp | 860 MiB |
| HG00735 paternal haplotype | 1 | 3.03 Gbp | 861 MiB |
| Plasmids | 72 556 | 7.30 Gbp | 2.1 GiB |
| Bacterial assemblies | 1 000 | 4.21 Gbp | 1.3 GiB |

**Table S2.** Command-line settings used for the set-operation benchmarks.

| Tool | Stage | Datasets | Settings |
| --- | --- | --- | --- |
| Sorted lists | build | human haplotypes | `sorted_lists build -k 31` |
| Sorted lists | build | plasmids, bacteria | `sorted_lists build -k 31 --strand both` |
| Sorted lists | set ops | all | `sorted_lists merge, intersect, difference` |
| KMC | build | human haplotypes | `kmc -k31 -m32 -t64 -ci1 -fm -b -r` |
| KMC | build | plasmids, bacteria | same as above, on preprocessed input including reverse complements |
| KMC tools | union | all | `kmc_tools -t<T> simple union -ocsum` |
| KMC tools | intersection | all | `kmc_tools -t<T> simple intersect -ocmin` |
| KMC tools | difference | all | `kmc_tools -t<T> simple kmers_subtract` |
| SBWT | build | human haplotypes | `sbwt build -k 31 -t 64 --in-memory` |
| SBWT | build | plasmids, bacteria | `sbwt build -k 31 -t 64 --mem-gb 32 --add-revcomp --dedup-batches --in-memory` |
| SBWT | set ops | all | `merge, intersect, difference -t<T> [--low-ram]` |

**Table S3.** Command templates used to construct the 661K SBWT index through a binomial merge pipeline. The `--low-ram` flag is used for merge levels at least 5, or as a fallback when a default merge fails.

| Step | Command template |
| --- | --- |
| Batch build, binomial pipeline | `sbwt build LIST -k 31 --mem-gb 48 --threads 32 --add-revcomp --dedup-batches --in-memory` |
| Merge, binomial pipeline | `sbwt merge --threads 32 [--low-ram]` |
| Direct build, in-memory | `sbwt build LIST -k 31 --mem-gb 48 --threads 32 --add-revcomp --dedup-batches --in-memory` |
| Direct build, disk-backed | `sbwt build LIST -k 31 --mem-gb 48 --threads 32 --add-revcomp --dedup-batches --temp-dir TMP` |

## Footnotes

1 These groups are marked in an additional bit vector during merge plan construction.

2 v0.1.0, Rust implementation available at https://github.com/LoreDepuydt/sbwt-set-operations

3 v0.1.0, Rust implementation available at https://github.com/LoreDepuydt/sorted-lists

